# Characterisation of acetogen formatotrophic potential using *E. limosum*

**DOI:** 10.1101/2022.11.02.514939

**Authors:** Jamin C. Wood, R. Axayacatl Gonzalez-Garcia, Dara Daygon, Gert Talbo, Manuel R. Plan, Esteban Marcellin, Bernardino Virdis

## Abstract

Formate is a promising energy carrier that could be used to transport renewable electricity. Some acetogenic bacteria, such as *Eubacterium limosum*, have the native ability to utilise formate as a sole substrate for growth, which has sparked interest in the biotechnology industry. However, formatotrophic metabolism in acetogens is poorly understood, and a systems-level characterization in continuous cultures is yet to be reported. Here we present the first steady-state dataset for *E. limosum* formatotrophic growth. At a defined dilution rate of 0.4 d^-1^, there was a high specific uptake rate of formate (280±56 mmol/gDCW/d), however, most carbon went to CO_2_ (150±11 mmol/gDCW/d). Compared to methylotrophic growth, protein differential expression data and intracellular metabolomics revealed several key features of formate metabolism. Upregulation of pta appears to be a futile attempt of cells to produce acetate as the major product. Instead, a cellular energy limitation resulted in the accumulation of intracellular pyruvate and upregulation of Pfl to convert formate to pyruvate. Therefore, metabolism is controlled, at least partially, at the protein expression level, an unusual feature for an acetogen. We anticipate that formate could be an important one-carbon substrate for acetogens to produce chemicals rich in pyruvate, a metabolite generally in low abundance during syngas growth.

## Introduction

Acetogens hold great promise for sustainable chemical and fuel production whilst closing the carbon cycle using feedstocks from renewable sources (Ljungdahl 2009). Acetogens can use reducing equivalents through intermediates such as hydrogen (H_2_), carbon monoxide (CO), formate and methanol. However, compared to synthesis gas fermentation (syngas, a mixture of CO and H_2_), as has been commercialised by LanzaTech using offgases from the steel industry, much less is known about the liquid C_1_ feedstock, formate (Cotton et al. 2020; Köpke and Simpson 2020).

Unlike the other liquid C_1_ feedstock methanol, formate can be efficiently produced directly from CO_2_ and renewable electricity without the need for hydrogen (*i.e*. Power-to-X), a technology that is at the pre-commercialisation stage (Spurgeon and Kumar 2018; Rabiee et al. 2021). Being a liquid, it avoids many of the transportation issues present with gaseous substrates, as well as fitting with existing supply chain and fermentation infrastructure (Cotton et al. 2020). It also can overcome key mass transfer limitations faced by gas fermentation, and have higher energy efficiencies, resulting in lower operational costs, *e.g*. for like cooling (Cotton et al. 2020). Formate could, in the future, become not only a biotechnology feedstock, but an energy carrier that avoids the pitfalls of hydrogen such as non-negligible supply chain leaks leading to additional costs and contribution to global warming as a result of its GWP over 100 years of 11 (that is, 11 times that of CO_2_) (Al-Breiki and Bicer 2020; Warwick et al. 2022).

Consequently, there have been numerous efforts to engineer formatotrophy into several organisms. For example, the synthetic reductive glycine pathway has been engineered into *E. coli* (Kim et al. 2020). The so-called FORCE pathway, an orthogonal chain elongation metabolism to biomass based on formyl-CoA, has also been engineered into *E. coli* (Chou et al. 2021). Despite these advances, native acetogen formatotrophy has the highest energy efficiency, which may be a critical metric when considering it as a Power-to-X technology (Wood et al. *In press;* Neuendorf et al. 2021). Utilising the native capabilities of acetogens such as *Eubacterium limosum, Butyribacterium methylotrophicum* and *Acetobacterium woodii*, would avoid the need for genetic engineering or building new gas fermentation infrastructure.

Formate as a substrate in the Wood-Ljungdahl Pathway (WLP) is similar to CO metabolism in that there is excess oxidation to CO_2_ in order to generate the required number of reducing equivalents. We have previously shown for *E. limosum* in batch culture that formate consumption results in acetate and CO_2_ production at a stoichiometric ratio of 1:1 (Wood et al. 2021). Interestingly, the observed native formatotrophic acetogen maximum growth rate was similar to that seen for the synthetic reduction glycine pathway in *E. coli* (*ca*. 0.1 h^-1^) (Kim et al. 2020).

Analytical quantification and metabolic modelling have advanced knowledge of acetogen metabolism, allowing for improved process economics and fermentation designs (Marcellin et al. 2016; Heffernan et al. 2022). Transcriptome and translatome data have been analysed for *E. limosum* (Song et al. 2017, 2018), and proteomic data for closely related *E. callanderi* (Kim et al. 2021). Some information has been published for the model acetogen, *A. woodii* (Neuendorf et al. 2021), yet no formatotrophic omics datasets exist for *E. limosum* to our knowledge. Since acetogen metabolism is generally acknowledged to be controlled post-transcription and translation, omics data, including metabolomics, may be key to revealing growth bottlenecks (Marcellin et al. 2016; Mahamkali et al. 2020; Heffernan et al. 2022). Moreover, in contrast to batch datasets, steady-state chemostat cultures offer better reproducibility across experiments, allowing greater insight into process optimisation (Heffernan et al. 2020).

To understand formatotrophic growth and limitations, in this study, we investigated steady-state C_1_ liquid formatotrophic fermentation in chemostat for *E. limosum*. Based on our previous batch observations showing distinct growth phases when grown on formate and methanol, here we hypothesised there would be limitations to continuous formate assimilation, perhaps similar to carbon monoxide toxicity leading to oscillatory growth observed in *Clostridium autoethanogenum* (Mahamkali et al. 2020). We tested this using differential analysis between formatotrophic and methylotrophic conditions. Intriguingly the differences are related to both protein expression and thermodynamics. Whilst growth was poor, unexpectedly, acetate was not the major product. Furthermore, we observed a biofilm formation, which would negatively impact scalable production. Anecdotally, adaptive laboratory evolution could be used to address this negative phenotype, particularly given some *E. limosum* strains are known to not form sticky polymers that support biofilms (Flaiz et al. 2021).

We understand there is currently a metabolic model under curation for *E. limosum*, and therefore differential proteomic and metabolomic data presented here may be a powerful tool to help facilitate construction of the model (Bae et al. 2021; Fackler et al. 2021).

## Methods/Experimental

### Bacterial Strain, Growth Medium, and Continuous Culture Conditions

*Eubacterium limosum* ATCC 8486 (*E. limosum*) was subject to more than 200 generations of Adaptive Laboratory Evolution under liquid C_1_ fermentation conditions (500 mM methanol, 100 mM formate). A single colony from this cell lineage was used in all experiments and stored as glycerol stocks at −80°C. This strain was cultivated anaerobically at 37 °C in a chemically defined phosphate buffered medium with liquid C_1_ feedstocks (Table 1). The medium contained per litre: 0.5 g MgCl_2_.6H_2_O, 0.5 g NaCl, 0.13 g CaCl_2_.2H_2_O, 0.75 g NaH_2_PO_4_, 2.05 g Na_2_HPO_4_, 0.25 g KH_2_PO_4_, 0.5 g KCl, 2.5 NH_4_Cl, 0.017 g FeCl_3_.6H_2_O, 0.5 g L-cysteine hydrochloride monohydrate, 1 mL of 1 g/L resazurin, 10 mL trace metal solution (TMS), 10 mL B-vitamin solution. The TMS contained per litre: 1.5 g nitrilotriacetic acid, 3 g MgSO_4_.7H_2_O, 0.5 g MnSO_4_.H_2_O, 1 g NaCl, 0.667 g FeSO_4_.7H_2_O, 0.2 g CoCl_2_.6H_2_O, 0.2 g ZnSO_4_.7H_2_O, 0.02 g CuCl_2_.2H_2_O, 0.014 g Al_2_(SO_4_)_3_.18H_2_O, 0.3 g H_3_BO_3_, 0.03 g Na_2_MoO_4_.2H_2_O, 0.028 g Na_2_SeO_3_.5H_2_O, 0.02 g NiCl_2_.6H_2_O and 0.02 g Na_2_WO_4_.2H_2_O. The B-vitamin solution contained per litre: 20 mg biotin, 20 mg folic acid, 10 mg pyridoxine hydrochloride, 50 mg thiamine-HCl, 50 mg riboflavin, 50 mg nicotinic acid, 50 mg calcium pantothenate, 50 mg vitamin B_12_, 50 mg 4-aminobenzoic acid and 50 mg thioctic acid. The Rnf complex in *E. limosum* is Na^+^ dependant, and therefore to maintain a consistent concentration across all experiments, NaCl was supplemented to medium which did not contain sodium formate as a substrate (*i.e*. chemostats growing on methanol). As a more oxidised substrate is required for methanol growth, gaseous CO_2_ was also provided.

**Table 1.** Summary of Eubacterium limosum fermentations. N = stirrer speed; BR = biological replicates; BC = biomass concentration. Methylotrophic data in the first row is provided for comparison from (Wood et al. 2022).

| Condition | Gas | N | BR | D | pH | BC | Acetate | Butyrate |
| --- | --- | --- | --- | --- | --- | --- | --- | --- |
|  |  | rpm | # | Day<sup>-1</sup> | Setpoint | gDCW/L | g/L | g/L |
| 100mM Methanol | 85% Ar, 15% CO<sub>2</sub> | 400 | 2 | 0.4 | 6.8 | 0.54±0.05 | 1.60±0.08 | 0.99±0.05 |
| 100 mM Formate | 100% Ar | 400 | 2 | 0.4 | 6.8 | 0.07±0.02 | 0.014±0.006 | 0.006±0.001 |

Steady-state conditions were reached in 0.5 L Multifors bioreactors (infors AG) controlled by EVE software at a working volume of 350 mL (magnetic marine impeller agitation). The system was equipped with peristaltic pumps, mass flow controllers (MFCs), pH and temperature sensors, and was connected to a Hiden HPR-20-QIC mass spectrometer (Hiden Analytical) for on-line off-gas analysis. Antifoam (Sigma 435503) was continuously added to the bioreactor using a syringe pump at 10 μL/h. Results presented here are after optical density, gas uptake/production rates, and acid/base addition rates were constant for at least five working volumes. Three technical replicate samples, spaced by one dilution volume, were collected per biological replicate.

### Experimental analysis and Quantification

#### Extracellular analysis

Optical density (OD) measurements were taken at 600 nm via a UV-Vis spectrophotometer (Thermo Fischer Scientific Genesys 10S UV-Vis Spectrophotometer, USA). A biomass formula of C_4_H_7_O_2_N_0.6_ and 0.32 gDCW/L/OD was used to convert OD to molar cell concentrations (Wood et al. 2021). Liquid samples for extracellular metabolomic analysis were collected, filtered and stored at −20 °C. Samples were analysed by high performance liquid chromatography using an Agilent 1200 HPLC System with Phenomenex Rezex RHM-Monosaccharide H+ column (7.8 × 300 mm, PN: OOH-0132-KO) and guard column (Phenomenex SecurityGuard Carbo-H, PN: AJO-4490). Analytes were eluted isocratically with 4 mM H_2_SO_4_ at 0.6 mL min^-1^ for 48 min, and column oven temperature of 65 °C. 30 μL of sample was injected and monitored using a UV detector (210 nm) and RID at positive polarity and 40 °C.

Bioreactor off-gas analysis was performed by an on-line mass spectrometer, monitoring the amounts of H_2_, Ar and CO_2_ at 2, 40 and 44 amu respectively. To achieve reliable off-gas analysis, a bypass line from the feed gas bottle was used as the calibration gas for each MS-cycle (*i.e*. ‘on-line’ calibration). Specific rates (mmol/gDCW/d) were calculated by accounting for the exact gas composition, bioreactor liquid working volume, feed gas flow rate, off-gas flow rate (based on a constant flow of inert Argon), the ideal gas molar volume, pH and CO_2_ dissolution equilibrium, and the steady-state biomass concentration.

As a quality check, gaseous samples were also taken for ‘off-line’ analysis using a Shimadzu 2014 GC, equipped with a ShinCarbon packed column (ST 80/100, 2mm ID, 1/8 OD Silico, Restek). H_2_ and Argon were detected by a thermal conductivity detector (TCD), while CO_2_ was measured using a flame ionization detector (FID).

#### Intracellular analysis

Quantitative proteome analysis was carried out using direct data-independent acquisition mass spectrometry approach (direct-DIA). Sampling, storage and sample preparation were performed based on a method previously developed for autotrophic growth of *C. autoethangenum*. Briefly, 5 ODs of culture was pelleted by immediate centrifugation (16,000g for 3 minutes at 4 °C) followed by washing with 1 mL PBS (Sigma P4417). A further round of centrifugation was used, with the supernatant then discarded and the remaining sample stored at −20 °C. Approximately 100 μg of protein was resuspended in 25 μL Milli-Q water and combined with 25 μL SDS lysis buffer (10% SDS, 100 mM Tris). Cells grown on formate required bead beating with a small amount of 0.1 mm diameter glass beads to lyse. To reduce protein disulphide bonds, dithiothreitol (DTT) was added to a final concentration of 20 mM before incubating at 70 °C for 1 hour, lodoacetamide (IAA) was added to a final concentration of 40 mM and incubated in the dark for 30 minutes to alkylate cysteine. The reaction was sonicated and 2.5 μL 12% phosphoric acid was added, before combining with 165 μL S-Trap binding buffer (90% methanol, 100 mM Tris (aq)). The sample was centrifuged for 8 minutes at 13,000 g, before adding to the S-Trap spin column, spun for 1 minute at 4000g and washed thrice with 150 μL S-Trap binding buffer. 1 μg of trypsin in 50 μL of 50 mM ammonium bicarbonate pH 8 was added to cover the top of the protein trap, incubating in a sealed bag overnight at 37 °C. Peptides were eluted into a collection tube with 40 μL increasing concentration of acetonitrile in 0.1% formic acid. Samples were spun dry and resuspended in 20 μL buffer A (0.1% formic acid (aq)) for injection to the mass spectrometer.

Mass spectrometry for proteomics analysis was performed using LC-MS/MS with a Thermo Fisher Scientific UHPLC system coupled to an Exploris 480 mass spectrometer. The electrospray voltage was 2.2 kV in positive-ion mode, and the ion transfer tube temperature was 250 °C. Full MS scans were acquired in the Orbitrap mass analyser over the range of m/z 340-1110 with a mass resolution of 120,000 (at m/z 200). The automatic gain control (AGC) target value was set at ‘Standard’, and the maximum accumulation time was ‘Auto’ for the MS/MS. The MS/MS ions were measured in 12 windows from mass 350-470, in 36 windows from mass 465-645 and 10 windows from mass 640-1100. Analysis of data were performed using Spectronaut against a reference proteome (UniProt ID UP000246246), with a Q-value cutoff of 0.05 applied to differential protein expressions. Locus tags correspond to (Song et al. 2017, 2018).

Intracellular metabolome analysis was based on the method previously developed for autotrophic growth of *C. autoethangenum* (Mahamkali et al (2020)). Briefly, 5 ODs of culture was pelleted by immediate centrifugation (16,000g for 3 minutes at 4 °C) followed by resuspension in chilled 50% acetonitrile to extract intracellular metabolites. Cell debris was removed by centrifugation with the supernatant stored at −80 °C. Samples were then freeze dried and resuspended in 90 μL 2% acetonitrile containing 5 μM azidothymidine standard. To remove lipophilic compounds (such as lipids, fatty acids, oil), the extract was fractionated by adding 250 uL of chloroform, before adding 410 μL MQ water, vortexing, and then separating the organic and polar phases by centrifugation. The polar fraction was cleaned through a spin column, freeze dried and resuspended with 2% acetonitrile. LC-MS/MS analysis was performed using a Shimadzu UHPLC System (Nexera X2) coupled to a Shimadu 8060 triple quadrupole (QqQ) mass spectrometer following (Espinosa et al. 2020; Mahamkali et al. 2020) with modifications and additions to the scheduled multiple reaction monitoring (sMRM) transitions. Chromatographic separation was performed on a Gemini NX-C18 column (3μm x 150mm x 2mm, PN: 00F4453B0, Phenomenex) with an ion-pairing buffer system consist of mobile phase A: 7.5mM tributylamine (pH 4.95 with acetic acid) and mobile phase B: acetonitrile. 5 μL and 10 μL of sample were injected to ensure measured intensities fell within the standard curve ranges.

Cell pellets were also used for PHB content analysis, following the method of (de Souza Pinto Lemgruber et al. 2019).

#### Thermodynamic and kinetic metabolic flux analysis

We evaluated reaction thermodynamic driving forces using the thermodynamic variability analysis model presented by Mahamkali et al. (2020). pH was assumed as 1 unit higher than the extracellular pH (Lindley et al. 1987). Reaction concentrations were constrained by measured values from intracellular metabolomics (or the lower limit of quantification, LLOQ). Carboxylate products (*i.e*., acetate, butyrate etc.) intracellular concentrations were limited according to between *ca*. 1 times higher, or 10^6^ times lower, than extracellularly, depending on possible transport mechanism (Lindley et al. 1987; Mahamkali et al. 2020). For formate substrate intracellular concentration, this was instead between 10 times higher, or 50 times lower, than extracellularly (Refer to Dataset A).

Beyond that, minimum concentrations were set to be 0.1 μM for metabolites, and 1 nM for dissolved gases unless directly measured or evaluated from Henry’s law (H_2_, CO_2_ and CO). CO_2_ total concentration, *c*, was calculated from partial pressure, *p*, as,

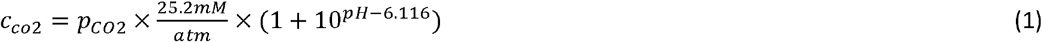

Maximum values were set to 1 mM for activated metabolites, and 10 mM for others.

A high-level estimate for reaction fluxes was determined as follows to estimate potential bottlenecks in metabolism. Considering metabolic flux (*J*) is controlled by protein concentration (E), Gibbs free energy of reaction (Δ*G*), saturation by substrates (*M*) with affinity (*K*), kinetic orders (*a*), and other sources of regulation (*U*), according to (Heffernan et al. 2022),

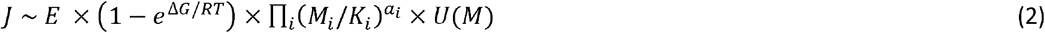

The enzyme and kinetics terms can be further simplified using Michaelis-Menten enzyme kinetics for a given substrate concentration (*S*) to give flux (*J*) as,

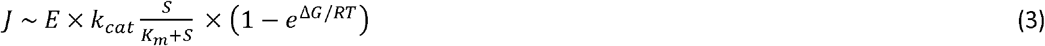

We can then make several assumptions to get likely order of magnitude estimates for *J* as follows.

- Our proteomic analysis can not determine absolute quantifications due to differences in mass spectrometer protein constituent signal response. However using a spiked protein analysis, (Valgepea et al. 2022) found a linear relationship between the log_10_ of mass spectrometer intensity and log_10_ anchor protein concentrations using a similar instrument setup to here. Since we can not be certain of the injected sample protein mass, we instead scaled protein concentrations by the pta enzyme, which Valgepea et al. (2022) showed was largely consistent across different gas fermentation conditions.
- Few *E. limosum* proteins have been assayed, and therefore instead we undertook BLAST analysis of key proteins against enzymes with known kinetic parameters, k_cat_ and K_m_. We then used this as verification data against deep learning models, where protein amino acid sequences were used to yield a complete set of kinetic parameters for the WLP central carbon metabolism (Kroll et al. 2021; Li et al. 2021). Where intracellular substrate concentrations, S, were unknown, instead k_cat_/K_m_ was taken as a proxy for kinetic effects.

## Results

### Steady state fermentation

Formatotrophic grown cells reached steady-state at a dilution rate (D) of 0.4 d^-1^ (specific growth rate of 0.017 h^-1^). The dilution rate was chosen as the highest achievable across formatotrophic and methylotrophic growth conditions, based on batch bottle growth rates. Despite the same amount of carbon being supplied to each condition, formate reached only a biomass concentration of 0.07±0.02 gDCW/L, less than 15% of methylotrophic growth. We note however, that there was significant biofilm formation in the formate condition, and so biomass concentration as reported here may be underestimated. As a result, substrate-specific uptake rates were much higher for the formate condition but potentially overestimated: 280±56 mmol/gDCW/d compared to 75±7.0 mmol/gDCW/d for methanol in the methanol/CO_2_ condition (**Figure 1A**).

**Figure 1.**
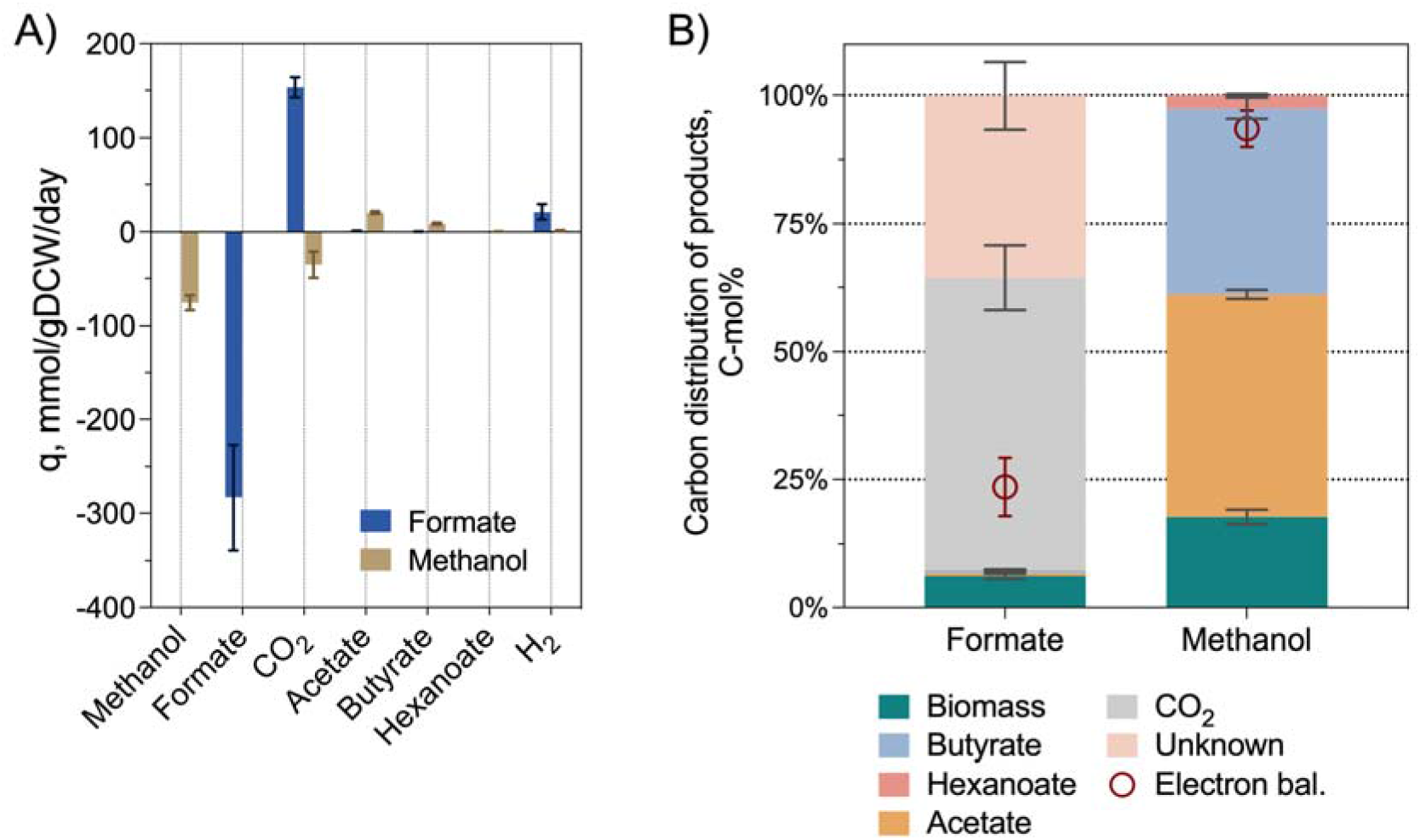
Characteristics of Eubacterium limosum in autotrophic chemostats, with conditions as summarised in Table 1. Methylotrophic data is provided for comparison from (Wood et al. 2022). (A) specific uptake and production rates. (B) carbon distribution of products. Values represent the average ± standard deviation between biological replicate triplicates.

Formatotrophic growth could only be achieved when no CO_2_ gas was supplied to the culture. Further, a formate titre of 2.0±0.18 g/L was observed at steady-state (**Table 1**), indicating the culture was not formate limited, but rather constrained by another parameter.

As expected, formate-grown cells showed significant CO_2_ evolution (150±11 mmol/gDCW/d) (a formate consumption to CO_2_ production ratio of 1.9±0.4 mol/mol). However, unexpectedly, the major non-CO_2_ product under formatotrophic growth was not acetate. Only trace amounts of acetate were detected throughout the entire fermentation (0.92±0.35 mmol/gDCW/d) (**Figure 1A**). Other typical *E. limosum* products such as butyrate (0.32±0.13 mmol/gDCW/d), hexanoate and butanol, were minor or below the detection limit (**Figure 1A**).

Hydrogen was produced in both cultures as a minor sink of electrons. Intriguingly, despite formate being the less reduced substrate, it produced more hydrogen at 21±7.8 mmol/gDCW/d compared to 1.1±0.56 mmol/gDCW/d for methanol/CO_2_ (**Figure 1A**). We noted in Wood *et al*. (2022) that formate seems to trigger hydrogen production, which is further reinforced with this data.

Overall, this gave a carbon and electron balance of 63±15% and 24±6% for formatotrophic growth (**Figure 1B**). These values suggest the production of an unknown product with a redox number of 5.0±2.8. Given none of the known products which branch from acetyl-CoA were detected (e.g. acetate, butyrate, hexanoate, butanol etc.), we hypothesised there may be a product downstream of pyruvate. Whilst no genes are identified in the *E. limosum* genome for PHB production, given the unknown product redox number, we analysed cell pellets for intracellular PHB content. Formate cultures showed 0.68±0.63% wPHB/wDCW. This is sufficiently low that the effect on carbon and electron balances is negligible. We also measured for 4-aminobutyrate (GABA) (potentially produced *via* partial TCA cycle) and 3-methyl-2-oxopentanoate (potentially produced *via* isoleucine biosynthesis) using HPLC analysis against standards. As neither of them could be found, we also looked for intermediates along the GABA pathway (2-ketoglutarate), and 3-methyl-2-oxopentanoate pathway (citraconate, 2-oxobutanoate). The analysis confirmed that none of these compounds were significantly detected in the samples, although the extracellular 2-ketoglutarate concentration was above the LLOQ for the formatotrophic condition, yet it was not detected for methylotrophic growth (data not shown).

### Intracellular conditions

Given the lack of success in identifying the missing product, we performed proteomics and metabolomics analyses to gain an understanding of formatotrophic *E. limosum* metabolism, and compared it to methanol metabolism (Wood et al. 2022). We identified 934 proteins that met the criteria for differential analysis between the formate and methanol/CO_2_ growth conditions (*i.e*. present in both samples and Q-value < 0.05). Proteome analysis detected all the key enzymes of the WLP, energy conservation, direct acetyl-CoA condensation pathway, carboxylate/alcohol production, central carbon metabolism, shikimate pathway, partial TCA cycle, isoleucine biosynthesis, and biofilm switches (**Figure 2** and **Figure 3**). Across these pathways, 31 intracellular metabolites were identified in both conditions. Coenzyme A was below the lower limit of quantification (LLOQ) for all conditions.

**Figure 2.**
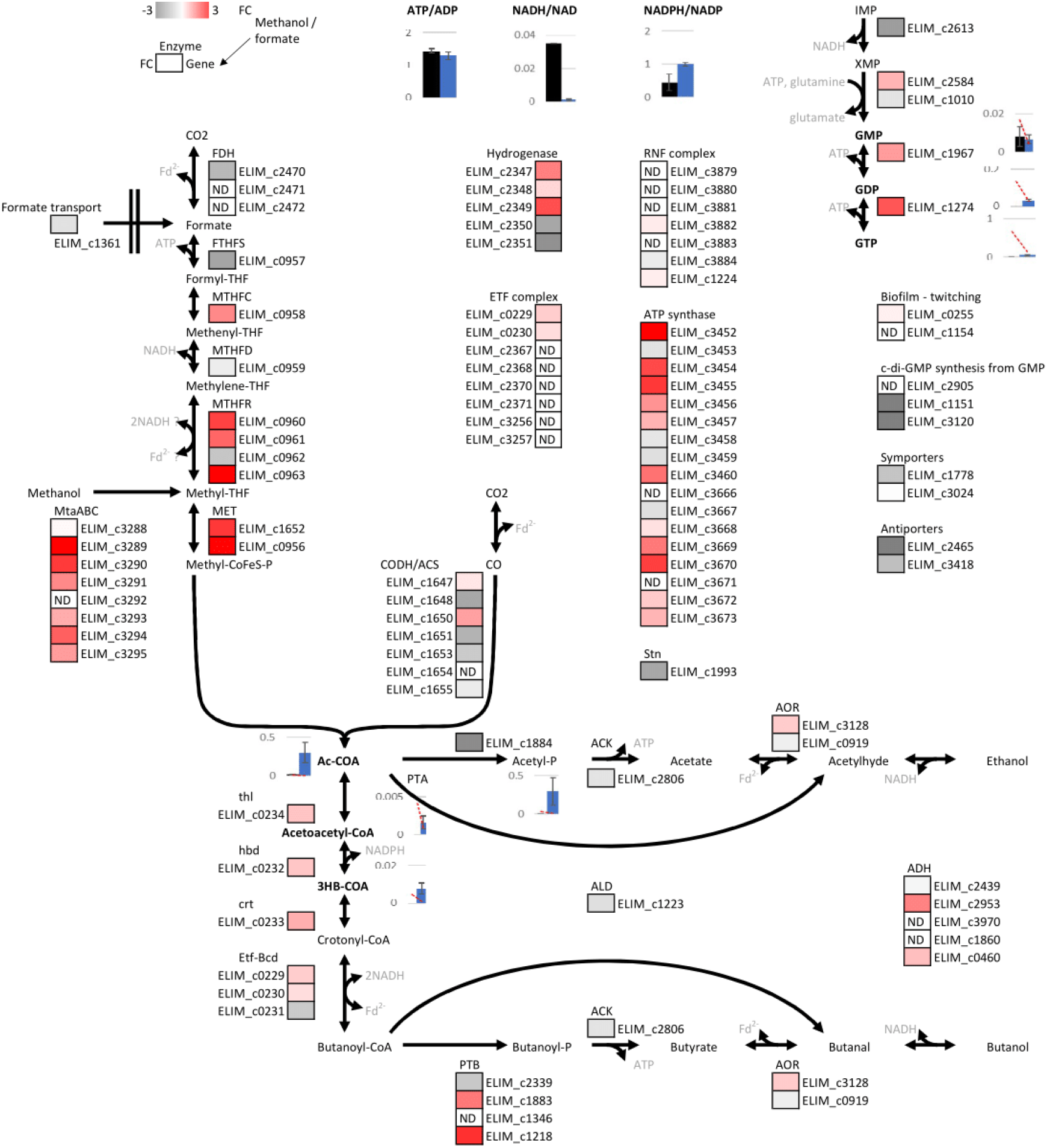
Schematic representation of Wood Liungdahl Pathway metabolism in E. limosum during C_1_ fermentation. Reactions are shown with substrates, products and redox mediators, however without stoichiometric balances along pathways. Enzymes and associated protein accession numbers (elim_cxxxx) are shown for each reaction, along with log_2_ fold changes comparing methylotrophic (Wood et al. 2022) and formatotrophic growth. Intracellular metabolites concentrations (μmol/gDCW) are shown for select metabolites as the average ± standard deviation of biological replicate triplicates. The methanol condition is in blue, and formate condition in black. The lower limit of quantification is shown as a red dashed line, which differs between the conditions due to the differences in cell pellet mass.

**Figure 3.**
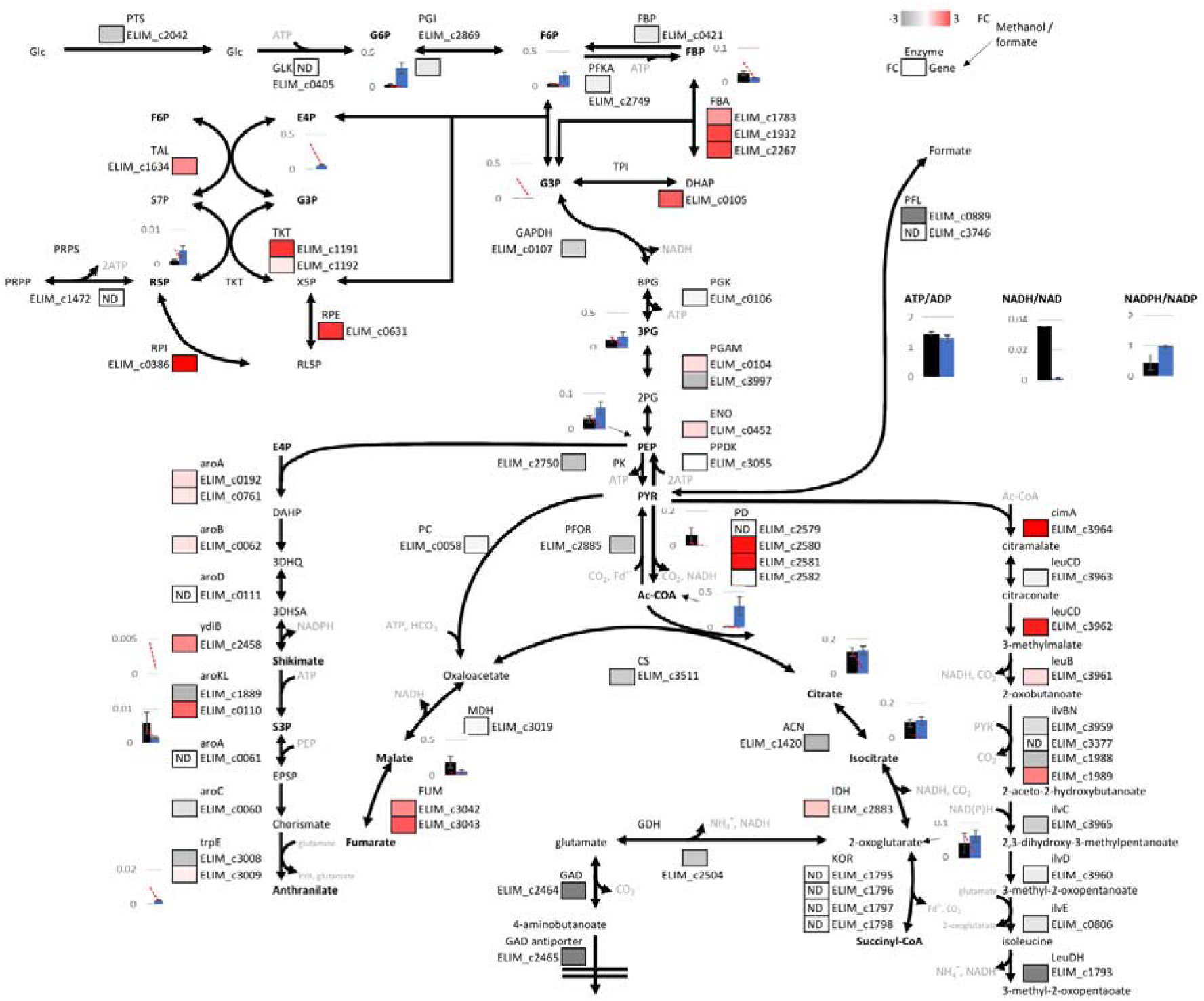
Schematic representation of central carbon metabolism metabolism in E. limosum during C_1_ fermentation. Reactions are shown with substrates, products and redox mediators, however without stoichiometric balances along pathways. Enzymes and associated protein accession numbers (elim_cxxxx) are shown for each reaction, along with log_2_ fold changes comparing methylotrophic (Wood et al. 2022) and formatotrophic growth. Intracellular metabolites concentrations (μmol/gDCW) are shown for select metabolites as the average ± standard deviation of biological replicate triplicates. The methanol condition is in blue, and formate condition in black. The lower limit of quantification is shown as a red dashed line, which differs between the conditions due to the differences in cell pellet mass.

Formate is assimilated via the formate dehydrogenase elim_c2470 and FTHFS elim_c0957 which have log_2_ fold changes of 1.7 and 2.1 and are respectively the 66^th^ and 8^th^ most ‘abundant’ proteins. Given the higher specific flux through the WLP for formate, it is not surprising there is upregulation generally (elim_c0959, 1648, 1651, 1655). That said, the log_2_ fold changes are mostly small and less than 2. The WLP proteins which are more regulated in the methanol/CO_2_ condition mostly relate to the thermodynamic difficulty in reversing methylene reductase (elim_c0960-63). Several of the WLP acs/codh complex were the most abundant in the formatotrophic proteome – elim_c1655, 1651. Yet, acetyl-CoA intracellular concentration was one order of magnitude lower (0.014±0.003 μmol/gDCW compared to 0.300±0.133 μmol/gDCW). For reference, *C. autoethanogenum* cells under syngas growth had an acetyl-CoA concentration of 0.358±0.140 μmol/gDCW (Mahamkali et al. 2020).

Strangely, mtaB (elim_c3293), part of the methyltransferase operon for methanol assimilation into the WLP was also one of the most ‘abundant’ proteins in the formatotrophic proteome. Song and colleagues showed no transcriptome fold change in mtaB between heterotrophic and autotrophic (H_2_/CO_2_) growth, and low relative expression, suggesting that our observation here shows formatotrophic and methylotrophic growth may share features not seen in other autotrophic conditions (Song et al. 2018).

When it comes to energy conservation, there was little change in etf proteins (elim_c0229, 0230), nor Rnf proteins detected (elim_c3882, 3884, 1224), yet ATPase was strongly downregulated in the formate condition (elim_c3452, 3454, 3455, 3670) with log_2_ fold changes greater than 2. ATP/ADP intracellular ratios were relatively consistent at 1.4±0.07 for the formate condition, and 1.3±0.11 for methanol/CO_2_, noting however the ATP/ADP/AMP pool size was one order of magnitude higher in the latter case. The sporomusa type Nfn complex was upregulated in the formate condition (elim_c1993 (Kremp et al. 2020)) with a log_2_ fold change of 2. The NADPH/NADP ratio was 0.40±0.25 and 1.0±0.056 for formate and methanol/CO_2_ respectively, with the same order of magnitude total pool size. The NADH/NAD ratios were 0.035 and 0.0011±0.0006 respectively, however the total pool size for methanol/CO_2_ was one order of magnitude higher. The formate condition ratio is similar to a typical syngas fermentation at NADH/NAD of 0.019 and NADPH/NADP of 0.276 (Mahamkali et al. 2020), which probably reflects formate and syngas operate the WLP and Nfn in the same direction, unlike methanol.

Considering pathways to produce carboxylates from acetyl-CoA (elim_c0231-0234, 2806, 2339) there is mostly little change in expression (log_2_ fold change < 1), reinforcing that this metabolism is regulated post-translation as acetate and butyrate were not major products under formatotrophic growth. Both acetoacetyl-CoA and 3-hydroxybutyryl-CoA were not detected, supporting the observation of no butyrate production. *E. limosum* codes for two different pta enzymes, one of which is significantly upregulated under methylotrophic growth (elim_c1218; log_2_ fold change 2.4), and the other significantly upregulated under formatotrophic growth (elim_c1884; log_2_ fold change 2.7). Despite the upregulation of elim_c1884, no acetyl-phosphate was detected for the formate condition, yet 0.294±0.177μmol/gDCW was detected under methanol/CO_2_ growth. This probably reflects the difference in acetyl-CoA/CoA driving force. Whilst cells did not produce alcohols under any condition, it is interesting to note identification of acetylating aldehyde dehydrogenase (elim_c1223), AOR (elim_c3128, 0919) and alcohol dehydrogenase (elim_c2439, 2953, 0460). None of these had log_2_ fold changes greater than 1.5 between the two conditions, yet it is worth noting elim_c3128 was in the top 5% of proteome ‘abundance’.

Carbon flux from the WLP is linked to pyruvate in glucogenesis via PFOR (elim_c2885) or alternatively Pfl (elim_c0889), the latter of which is upregulated with a log_2_ fold change of 5.4 for formatotrophic growth. This is the 9^th^ most upregulated formatotrophic gene indicating that cells may utilise formate directly to synthesise pyruvate, rather than oxidise it to generate ferredoxin and CO_2_ for use by PFOR. Pyruvate was not detected in the methanol/CO_2_ condition (LLOQ 0.0016 μmol/gDCW), yet it was 0.060±0.044 μmol/gDCW in formate growth.

The remainder of the glucogenesis pathway (elim_c3055, 0452, 3997, 0104, 0106, 0107, 0421, 2869) all have log_2_ fold changes less than 1.5. Two notable exceptions are TPI (elim_c0105) and FBA (elim_1932, 2267) which are downregulated for formatotrophic growth (log_2_ *ca*. 2), along with parts of the pentose phosphate pathway (elim_c0386, 0631, 1191) (log_2_ fold change *ca*. 2-3). Of all the measured intermediates, only FBP was higher under the formate growth condition (yet still below the LLOQ). This seems counterintuitive based on protein expression, unless FBP flux also goes elsewhere for methylotrophic growth.

Branching off the glucogenesis and pentose phosphate pathways, the shikimate pathway proteins all have log_2_ fold changes less than 1.5, with the sole exception of aroKL (elim_c0110) being downregulated in formatotrophic growth with a log_2_ fold change of 1.8. PEP and E4P, which enter the shikimate pathway, have lower concentrations for the formate condition, compared to methylotrophic growth (0.028±0.009 μmol/gDCW v. 0.060±0.018 μmol/gDCW, and N.D. v 0.058±0.012 μmol/gDCW). Downstream of the shikimate pathway, anthranilate was not detected in the former condition, and was 0.002±0.0004 μmol/gDCW in the latter case.

The partial TCA cycle, and isoleucine biosynthesis pathways both have three of the most upregulated proteins under formatotrophic growth (11^th^, 28^th^ and 43^rd^). GAD (elim_c2464) and its antiporter (elim_c2465) have log_2_ fold changes of 5.0 and 3.6 respectively, suggesting 4-aminobutyrate (GABA) may be an extracellular metabolite. Similarly, LeuDH (elim_c1793) has a log_2_ fold change of 3.9, suggesting 3-methyl-2-oxopentanoate may also be an extracellular metabolite. ACN (elim_c1420), which reversibly converts citrate and isocitrate was upregulated in the formate condition by a log_2_ fold change of 1.7. Yet, intracellular metabolite concentrations were the same in the two conditions (0.127±0.027 μmol/gDCW v. 0.136±0.026 μmol/gDCW, and 0.093±0.017 μmol/gDCW v 0.103±0.022 μmol/gDCW). Interestingly FUM (elim_c3042, 3043) were both downregulated with a log_2_ fold change of 1.4 and 2.0 respectively, which seems to correlate with the substrate, malate, having a higher intracellular concentration for formate (0.186±0.077 μmol/gDCW) compared to methanol/CO_2_ (0.056±0.018 μmol/gDCW). Isoleucine biosynthesis cimA (elim_c3964) and leuCD (elim_c3962) were downregulated by log_2_ fold changes of 4.3 and 2.6 respectively. The remainder of the partial TCA cycle and isoleucine biosynthesis pathways showed no significant change in regulation.

We found several significantly ‘abundant’ proteins that have not been annotated. These include, for example, elim_c2694 (log_2_ fold change 7.8, 22^nd^ most ‘abundant’), yet we note that it had no significant transcriptome fold changes (log_2_ < 1.5) between heterotrophic and autotrophic growth, indicating it may be specific to C_1_ liquid metabolism (Song et al. 2018). Further, there are also upregulated proteins for which the annotation is ambiguous. For example, elim_c2880 is the most upregulated protein with a log_2_ fold change of 10.4, as well as being the 5^th^ most ‘abundant’ protein. It is annotated as a BadM/Rrf2 transcriptional regulator, associated with benzoyl-CoA degradation, even though benzoate was not present in the medium. We do note BadL, used in PABA metabolism, is part of this operon although not annotated in *E. limosum* (Vandrisse and Escalante-Semerena 2018). Therefore, this could *potentially* mean there is a PABA nutrient limitation (noting that PABA was not detected intracellularly in either condition). Similarly, elim_c1937 is a folate transport upregulated for formatotrophic growth (log_2_ fold change of 3.8 compared to methylotrophic growth), indicating a potential THF limitation.

## Discussion

### Metabolic model and redox balances

Thus far we have presented ‘omics data for formatotrophic *E. limosum* cultures, with differential analysis against methylotrophic growth. Whilst valuable conclusions can be drawn, the data takes on further meaning when integrated together in a complete kinetic model. Since such a model does not exist for *E. limosum*, we have instead undertaken thermodynamic calculations and various kinetics assumptions to identify several potential metabolic bottlenecks (refer to materials and methods).

Unlike NADH/NAD and NADPH/NADP, we could not measure ferredoxin. However, upper and lower bounds on the ferredoxin state could be calculated. From the Stn, Fd^2-^/Fd must be > 0.075. Using Henry’s law, the dissolved hydrogen must be *ca*. 0.6 μM and so the hydrogenase suggests Fd^2-^/Fd must be > 2.4. With the Rnf/ATPase complex operating in the forward direction for formate metabolism (*i.e*., generating ATP), Fd^2-^/Fd should be > 1.5. For PFOR to generate pyruvate, and assuming a CoA concentration of <43 μM (LLOQ), Fd ^2-^/Fd must be <60. However, considering the predicted kinetic parameters, it is likely that the Fd^2-^/Fd ratio is on the lower end (*ca*. 4) to generate sufficient flux through the mthfr, codh and fdh. However, given that ferredoxin is mainly used in the WLP and pyruvate synthesis, this wide range of possible values (2.4 < Fd^2-^/Fd < 60) does not provide further clues as to identification of the unknown product.

**Figure 4** illustrates how methanol and formate metabolism differ in the WLP by reversing key enzymes. The hydrogenase thermodynamic driving force is much larger under formatotrophic growth, which may explain why we observed higher hydrogen evolution. The methanol condition has an acetyl-CoA/CoA ratio three orders of magnitude higher than the formate condition, yet we calculated a similar thermodynamic driving force. This is because, in the latter condition, a much lower acetate concentration is possible, hence why it is in fact possible for pta to reverse under formatotrophic growth. We calculated this acetate limit to be in the *ca*. 1mM range (using eQuilibrator (Flamholz et al. 2012) with acetyl-P = 12.5μM, ATP/ADP = 1.5, pH = 7.9, pMg = 3.0, ionic strength = 0.25). This confirms, that despite significant upregulation of the pta (elim_c1884), acetate production is limited by post-translation effects. Additionally, we cannot identify how thl is feasible to satisfy butyrate flux, without the acetoacetyl-CoA concentration being below our assumed minimum value of 0.1 μM, which would surely impose kinetic limitations. Thus, the model confirms our suspicion that acetyl-CoA must be diverted elsewhere (*i.e*., towards pyruvate).

**Figure 4.**
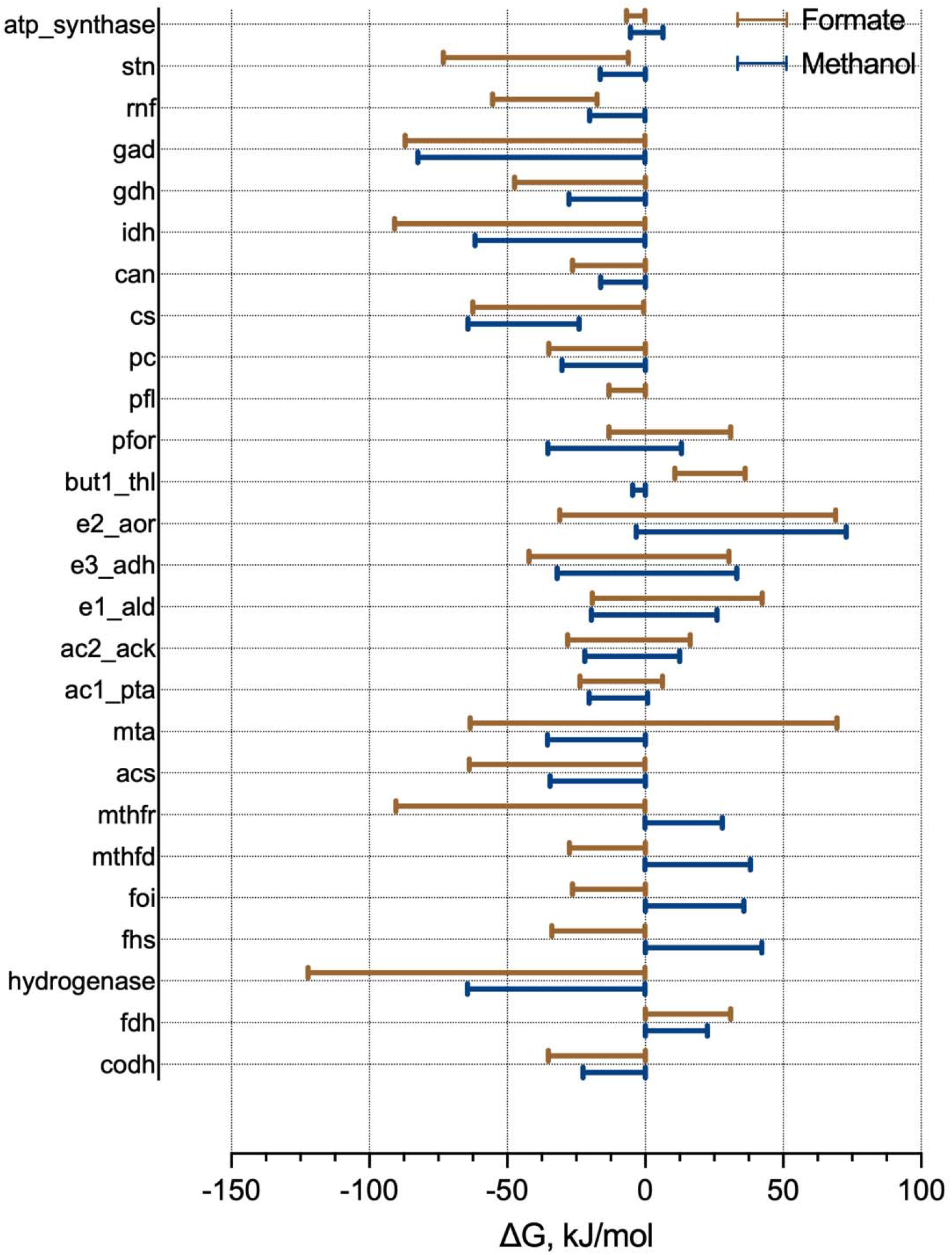
tMFA of E. limosum showing maximum allowable range of Gibbs free energy for respective reactions of the methanol/CO_2_ (blue) and formate (brown) conditions using thermodynamic variability analysis. Reaction directions are in the anabolic direction from CO_2_ on Figure 2, Figure 3. Rnf, PFOR, hydrogenase and stn are all calculated in the ferredoxin-consuming direction, and ATP synthase in the ATP-forming direction. Pfl is not shown for methylotrophic growth as it was not detected in the proteome.

Interestingly, the upregulated Pfl is probably the major source of pyruvate, considering thermodynamics (**Figure 4**), protein abundance and k_cat_/K_m_ (**Table 2**). Yet, the model indicates PFOR may in fact operate in the pyruvate consuming direction during formatotrophic growth, causing a futile metabolic cycling of pyruvate. Considering thermodynamics, protein ‘abundance’ and k_cat_/K_m_ (**Table 2, Figure 4**), these reinforce a mix of pre- and post-translation effects control metabolism and may explain the lack of production of GABA (no feasible export) and 3-methyl-2-oxopentanoate (kinetic bottleneck).

**Table 2.**
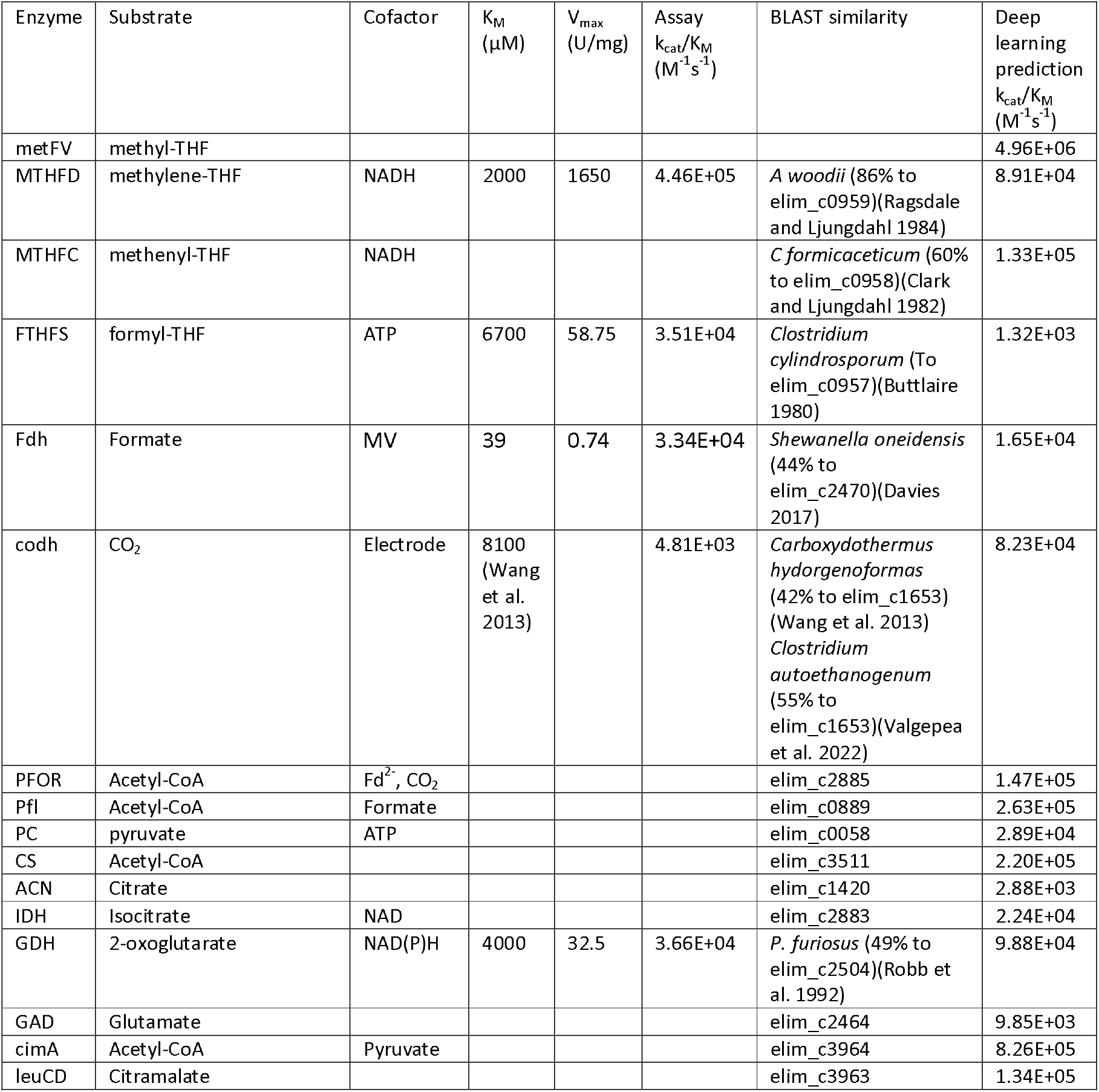
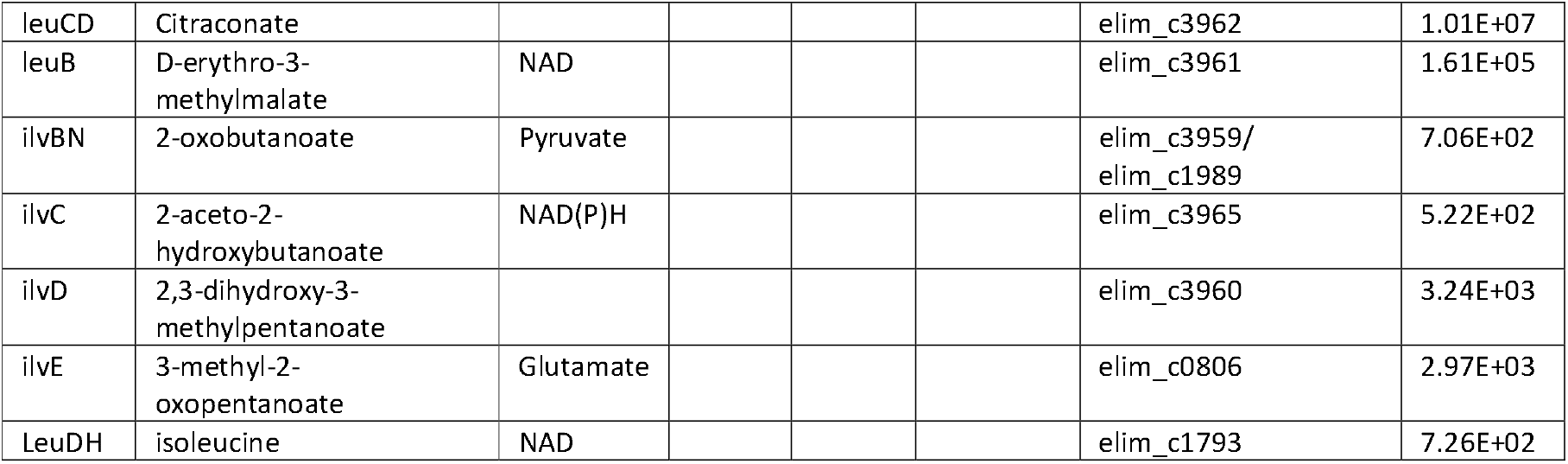
Kinetic parameters of enzymes with highest, to our knowledge, BLAST similarity to E. limosum enzymes. Deep learning models have been used as a comparison, and for all remaining pathways.

Given the Stn expression, NADPH likely has greater flux in the formate condition than methanol. This is interesting because the NADPH/NADP is lower for formate. This could be a clue for product identification, as production may be tied to the need to regenerate NADP. However, it is noted this would come at an ATP cost, as ferredoxin is diverted away from Rnf/ATPase chemiosmotic ATP generation, instead towards Stn to generate NADPH.

### Cellular stress and pyruvate

Formate metabolism in acetogens appears partially controlled by protein expression in response to cellular state. For example, since formatotrophic growth compared to methanol/CO_2_ has a much lower CO_2_ driving force, it makes sense that most pyruvate flux would be through heavily upregulated Pfl, despite PFOR being the normal mechanism for pyruvate synthesis (Neuendorf et al. 2021). Given the low acetyl-CoA/pyruvate ratio, Pfl would require a high formate concentration as driving force, which may explain why a formate titre existed at steady-state. Previously, researchers have suggested an increased pyruvate concentration is regulated by PPDK activity and ATP availability, causing a drop in glucogenic intermediates for *E. limosum* (Lebloas et al. 1996). Similarly, we found that formatotrophic growth had lower glucogenic intermediates and no change in PPDK levels (elim_c3055) compared to methylotrophic growth. Specifically, we can see that the PEP / pyruvate ratio is respectively 1.3±0.61 and >39±12 (adopting LLOQ for pyruvate where not detected). This means PPDK activity is being limited by ATP availability, and so increasing pyruvate concentration and hence the required formate driving force. This associated energy shortage allows redirection of carbon (Lebloas et al. 1996).

We can see this redirection of carbon in the lack of acetate production, and futile upregulation of the pta enzyme. Given that metabolites on the butyrate production pathway were not detected, this supports the idea that a significant amount of carbon goes through pyruvate.

Of all measured intracellular metabolites, most were relatively similar for both methanol and formate conditions. However, several which were much lower or below LLOQ for formate only. Some of these are on the glucogenesis pathway, which is controlled by ATP availability discussed above. We found that pools of NAD, ATP and acetyl-CoA to be one order of magnitude lower for formatotrophic growth. This is also significantly lower than typical acetogens, despite the NADH/NAD, ATP/ADP ratios being conserved relative to other conditions (Mahamkali et al. 2020). Clearly, these ratios are tightly controlled, despite the lower ATP gain from growth, but the overall pool size is not.

It is hard to understand why ATP, NAD and CoA pool sizes are smaller for formatotrophic growth compared to methylotrophic growth based on the proteomics data given that two proteins involved in NAD biosynthesis (elim_c1173, 3031) were upregulated. The remaining two (elim_c1197, 1880) were not detected. This includes elim_c1880, which is used to synthesise NADP from NAD, despite NADP(H) being detected as a metabolite. In CoA biosynthesis, only one (elim_c1486) of the four proteins were identified in the formate condition, compared to three (elim_c1486, 3523, 2435) in the methanol condition. The lack of protein detection may correlate with the reduced ability of cells to synthesise NAD and CoA. GTP is required for AMP synthesis (elim_c1406, 2669), and since the GMP flux is not directed to GTP in formatotrophic growth, this may affect the ATP pool size. Some researchers have found pool size can be reduced by cellular stress conditions, such as pH and temperature (Chohnan et al. 1998).

### Formate itself may be cause of stress

Carboxylate transport in acetogens is possible *via* passive diffusion, symport of anion with a proton, transport of anion *via* a uniport and ATP-consuming ABC transporter (Mahamkali et al. 2020). Given we are already energetically limited, we can discount the latter process. An anion uniport (e.g. elim_c1361 log_2_ fold change of 0.7) must balance the charge with independent proton uptake, which could be *via* an ATPase, thus assisting with the energetic requirements. This would yield a lower intracellular formate concentration driving force (1 mM; Dataset A), however, our model predicts the resulting intracellular formate driving force is still sufficient for cell function.

Given acid was added throughout the experiment to maintain extracellular pH, there must have been a net alkalinisation of the cell, which may be the cause of cellular stress. This would lead to a more negative membrane potential that might be balanced by importing protons through an ATPase. This could theoretically improve the overall ATP gain. It is important to note *E. limosum* ATPases are generally thought to be sodium-dependent and we also did not find a significant upregulation of any ATPase under formatotrophic growth (Song et al. 2017; Kang et al. 2020). Whilst there was clearly an ATP limitation, the ATP balance itself thus remains unclear.

Additionally, formate metabolism is known to lead to a loss of the acetyl-phosphate pool via formyl-CoA (and hence ATP) (Sly and Stadtman 1963). However, we have not identified the culprit CoA-hydrolase enzyme in the proteome. Overall, we speculate that cellular stress caused an energy shortage, increased pyruvate concentration, which then allowed redirection of carbon.

### What does AOR do?

We found high ‘abundance’ of AOR, with little difference between the two conditions. Previous researchers have noted that acetogens are regulated post-translationally (Marcellin et al. 2016; Mahamkali et al. 2020; Heffernan et al. 2022), and we believe this could be the case for AOR. Whilst the enzyme is known to be promiscuous, AOR is generally acknowledged not to have C_1_ specificity for formate (Bertsch and Müller 2015). The same can be said for adh. The kinetic deep learning model, however, suggests formate consumption may be possible (as k_cat_ and K_m_ are similar to C_2_ and C_4_ substrates, but adh k_cat_ is one order of magnitude lower for C_1_; data not shown), but would result in only *ca*. 0.2 μM accumulation of methanol, which is below our limit of detection. In principle, it is possible, which may explain our previous observation of methanol production from formate during early growth stages, presumably when there is a highly reduced redox pool (Wood et al. 2021).

### CO_2_ must be maintained low to allow growth in chemostat

We found cells were unable to grow formatotrophically when 15% CO_2_ gas was present in the headspace. Growth was restored, and steady state was reached when the gas feed contained zero CO_2_. If this requirement was thermodynamics related, the bottleneck reaction would therefore produce CO_2_. All else being equal, increasing the CO_2_ concentration means the formate driving force must increase, and there must be an upper limit to intracellular formate concentration in terms of toxicity.

### Biofilm formation

Two *E. limosum* cyclic-di-GMP-l riboswitches are known to regulate virulence, motility, quorum sensing and biofilm (Song et al. 2017). Here, under formatotrophic growth we found upregulation of two proteins which synthesise cyclic-di-GMP from GMP – elim_c1151, 3120 (log_2_ fold change of 6.2 and 3.1 respectively). Intracellular GMP concentrations were similar for both formatotrophic growth and methylotrophic growth at 0.008±0.005 μmol/gDCW and 0.007±0.002 μmol/gDCW, respectively. GMP can also be used for GDP and GTP synthesis and interestingly these proteins (elim_c1967, 1274) are downregulated for formatotrophic growth (log_2_ fold change 1.2 and 2 respectively). Additionally, unlike the methanol condition, no GDP or GTP were detected under formate growth, reinforcing that GMP flux went elsewhere. Therefore, we confirm cyclic-di-GMP has a correlation with biofilm formation strength.

We also note GlmU (elim_c1471), which produces a precursor for the adhesion of biofilms in some bacteria, was upregulated in the formate condition (log_2_ fold change of 1.1) (Burton et al. 2006). Interestingly GlmU transcriptome shows upregulation in heterotrophic growth, compared to autotrophic, which matches the past *E. limosum* biofilm formation observations (Song et al. 2018). LuxR (elim_c1099) is a transcriptional regulator known to affect biofilm formation through quorum sensing, and here it was upregulated in formatotrophic growth (log_2_ fold change of 2.4) (Chen and Xie 2011). There was little difference in one identified protein related to cell motility (twitching) (elim_c0255).

From the ‘omics data, biofilm formation in *E. limosum* formatotrophic growth is most likely caused by cyclic-di-GMP production, adhesive compounds and/or quorum sensing. Inhibitors for adhesive compounds produced by GlmU include N-acetyl glucosamine-1-phosphate and iodoacetamide (Chen et al. 2015). The latter reacts with cysteine in our medium, and so is not an option here. Anthranilate, a tryptophan metabolite, is known to reduce intracellular cyclic-di-GMP, which should reduce biofilm formation (Li et al. 2017). Other metabolites on the tryptophan pathway, including indole and D-tryptophan have also been shown to influence biofilm formation. However what works for one species, can even enhance formation in other species (Li et al. 2017).

Interestingly, we did not detect anthranilate in the formate condition, but it was 0.002±0.0004 μmol/gDCW under methylotrophic growth. Anthranilate is produced from S3P, which was three times higher in the formate condition (0.006±0.003 μmol/gDCW versus 0.002±0.0003 μmol/gDCW for methanol). This trend suggests that there is more flux to anthranilate, but its concentration is being kept low, perhaps to regulate intracellular cyclic-di-GMP. Hence more biofilm formed in the formate condition.

To investigate this relationship, we supplemented the medium with 0.8 mM anthranilate in a culture with a pre-formed biofilm. Li and colleagues noted approximately this amount of anthranilate reduced pre-formed biofilm coverage by up to 30% for three different species of bacteria (Li et al. 2017). However, to our surprise, this concentration of anthranilate had no visual effect on the formed biofilm (data not shown). It is possible anthranilate may only prevent biofilm formation in *E. limosum*, which we have not tested here. Alternatively, other biofilm inhibitors may need to be tried, such as N-acetyl glucosamine-1-phosphate to prevent adhesive compound synthesis (Chen et al. 2015).

## Conclusion

We have established a baseline formatotrophic steady-state dataset for *E. limosum* covering phenomics, proteomics and metabolomics. Cells had a high formate-specific uptake rate. However, unexpectedly they did not produce acetate as a major product. There is evidence of a cellular energy limitation, which resulted in an accumulation of pyruvate intracellularly, and depletion of the total CoA and NAD pools. Together, these redirected carbon flux away from acetate, despite upregulation of pta, and towards products downstream of pyruvate. We contend this state of cellular stress ultimately relates to formate itself as a substrate, which cells attempt to overcome through protein expression, for example, upregulation of Pfl. This is interesting because acetogen metabolism is usually controlled post-translation.

Despite having high energy efficiency, there are significant challenges to overcome for *E. limosum* formatotrophy to be a scalable technology. Firstly, limiting biofilm formation, either through medium supplementation or adaptive laboratory evolution, will be important to reduce operational costs. Formatotrophy also needs to overcome poor yields and limit excess CO_2_ production, necessitating another substrate such as hydrogen or methanol. Adopting either of those would partially defeat the purpose of using formate (*i.e*., liquid feedstock that can be directly synthesised from CO_2_, fits with existing infrastructure and has no mass transfer limitations). Throughout this work, we have identified exciting aspects of formate metabolism that could have significance for acetogen applications, which include more targeted products downstream of pyruvate.

## Abbreviations

General

D: Dilution rate
LLOQ: Lower Limit of Quantification
OD: Optical Density
WLP: Wood-Ljungdahl Pathway

Compounds

2PG: 2-phosphoglycerate
3DHQ: 3-dehydroquinate
3DHSA: 3-dehydroshikimate
3PG: 3-phosphoglycerate
Ac-CoA: acetyl-CoA
BPG: 1,3-biphosphoglycerate
DAHP: 3-deoxy-arabino-heptulonate 7-phosphate
DHAP: Dihydroxy acetone phosphate
E4P: Erythrose 4-phosphate
EPSP: 3-enolpyruvyl-shikimate 5-phosphate
F6P: Fructose 6-hosphate
FBP: Fructose 1,6-biphosphate
G3P: Glyceraldehyde 3-phosphate
G6P: Glucose 6-phosphate
Glc: Glucose
PEP: Phosphoenolpyruvate
PRPP: 5-phosphoribosyl diphosphate
PYR: Pyruvate
R5P: Ribose 5-phosphate
RL5P: Ribulose 5-phosphate
S3P: Shikimate 3-phosphate
S7P: Sedoheptulose 7-phosphate
THF: tetrahydrofolate
X5P: Xylulose 5-phosphate

Enzymes

ALD: aldehyde dehydrogenase
ACK: acetate kinase
ACN: anonitase
ACS: acetyl-CoA synthase
ADH: alcohol dehydrogenase
AOR: aldehyde:ferredoxin oxidoreductase
aroA: DAHP synthase
aroA: 3-phosphoshikimate 1-carboxyvinyltransferase
aroB: 3-dehydroquinate synthase
aroC: chorismate synthase
aroD: 3-dehydroquinate dehydratase
aroKL: shikimate kinase
BK: butyrate kinase
cimA: citramalate synthase
CODH: CO dehydrogenase
crt: crotonase
CS: citrate synthase
ENO: enolase
Etf-Bcd: butyryl-CoA dehydrogenase
FBA: Fructose biphosphate aldose
FBP: fructose 1
FDH: formate dehydrogenase
FTHFS: formyl-THF synthetase
FUM: fumarase
GAD: glutamate decarboxylase
GAPDH: glyceraldehyde 3-phosphate dehydrogenase
GDH: glutamate dehydrogenase
GLK: Glucokinase
hbd: 3-hydroxybutyryl-CoA dehydrogenase
IDH: isocitrate dehydrogenase
ilvBN: pyruvate:2-oxobutanoate acetaldehydetransferase
ilvC: 2-aceto-2hydroxy-butanoate:NADP+ oxidoreductase
ilvD: dihydroxy acid dehydratase
ilvE: branched chain amino acid aminotransferase
KOR: oxoglutarate ferredoxin oxidoreductase
leuB: methylmalate dehydrogenase
leuCD: methylmalate hydrolase
LeuDH: leucine dehydrogenase
MDH: malate dehydrogenase
MtaABC: methanol dependent methyl transferase
MTHFC: methenyl-THF cylcohydrolase
MTHFD: methylene-THF dehydrogenase
MTHFR: methyltransferase/methylene-THF reductase
PC: pyruvate carboxylase
PD: pyruvate dehydrogenase
PFKA: Phosphofructose kinase
PFL: pyruvate formate ligase
PFOR: pyruvate ferredoxin oxidoreductase
PGAM: phosphoglycerate mutase
PGI: Glucose 6-phosphate isomerase
PGK: phosphoglycerate kinase
PK: pyruvate kinase
PPDK: phosphoenolpyruvate synthetase
PRPS: Ribose phosphat epyrophosphokinase
PTA: phosphotransacetylase
PTB: phosphotransbutyrylase
PTS: phosphotransferase system
RPE: Ribulose phosphate epimerase
RPI: Ribose phosphate isomerase
Stn: Sporomusa type Nfn transhydrogenase
TAL: Transaldolase
thl: thiolase
TKT: Transketolase
TPI: Trisephosphate isomerase
trpE: anthranilate synthase
ydiB: shikimate dehydrogenase

## Availability of data and materials

The datasets used and/or analysed during the current study are available from the corresponding author on reasonable request.

## Authors’ contributions

JCW designed and conducted experiments, interpreted results and was a major contributor in writing the manuscript. RAGG jointly conducted experiments. DD performed intracellular metabolomics analysis. GT performed proteomics analysis. MRP performed extracellular metabolomics and PHB analysis. EM and BV conceived experiments and provided substantiative manuscript review. All authors read and approved the final manuscript.

## Acknowledgements

Not applicable

